# Genome-wide variant analysis reveals divergent genetic signatures in *Aedes aegypti* and its morphological variant *Aedes aegypti* var. *luciensis*

**DOI:** 10.1101/2024.10.22.619611

**Authors:** Bhavna Gupta, G Navaneetha Pandiyan, Melveettil Kishor Sumitha, Rajaiah Paramasivan, Mariapillai Kalimuthu

## Abstract

In this study, we analysed the genetic diversity between the two morphotypes of *Aedes aegypti* by identifying and characterizing genomic variants from low-coverage resequencing data. Four samples from each morphotype were sequenced, and high-quality variants were identified. Variants from the four samples of each morphotype were combined, considering only those present in all samples (missing variants were excluded). This resulted in 7,181 variants in the normal type and 4,513 in the variant type, with only 1,180 shared variants identified between the two morphotypes. Most variants in both types were found in non-coding regions. Chromosomal distribution showed comparable variant frequencies between morphotypes, with similar transition/transversion ratios. The neighbor-joining tree indicated distinct genetic clusters for each morphotype, likely driven by different demographic histories. We have not obtained much data on variations in the coding regions, probably due to low sequencing coverage. However, the significant genetic differentiation between the two morphotypes indicates important evolutionary processes at play. Understanding these variations is crucial from an epidemiological perspective, as they may lead to differences in vector competence for pathogens, potentially affecting transmission dynamics and disease risk.

## Introduction

*Ae. aegypti* mosquitoes pose a significant public health concern due to its widespread distribution and ability to transmit multiple arboviral diseases. The global consequences of *Aedes*-borne diseases, such as dengue, Zika, and chikungunya, have escalated, placing a substantial burden on public health systems (Roiz et al. 2024). This species thrives in urban environments, making it a formidable vector for arboviruses.

Mosquitoes encompass a vast array of vectors for various pathogens, often including numerous sibling or cryptic species (Zheng 2020). This phenomenon is particularly evident among Anopheles species, the primary vectors of malaria (Gunathilaka and Karunaraj 2015; Khan et al. 2022). Among these cryptic species, some act as vectors for pathogens, while others are refractory. Additionally, these cryptic species tend to exhibit specificity to particular geographical regions and breeding conditions, complicating vector ecology and disease transmission dynamics (Walton et al. 1999). In contrast, the situation with *Ae. aegypti* is comparatively straightforward, with only two well-known forms: *Ae. aegypti formosus*, confined to African forests, and *Aedes aegypti aegypti*, found globally (Mattingly 1957; Christophers 1960). However, numerous morphological varieties exist beyond these forms (Diouf et al. 2020; Kumar et al. 2022).

Although *Ae. aegypti* is classified as a single species worldwide, extensive genetic variations have been reported within different countries (Failloux et al. 2002; Gloria-Soria et al. 2016; Lv et al. 2020; Zhang et al. 2022; Sumitha et al. 2023; Zhao et al. 2023; Acharya and Singh 2024; Joyce et al. 2024). Beyond genetic differences, populations from various locations exhibit varying levels of competence and transmission efficiency for pathogens. For examples, populations from different regions of Mexico, Brazil showed significant variations in the competence and transmission efficiency for pathogens (Bennett et al. 2002; Garcia-Luna et al. 2018). Similar type of variation in the competence and susceptibility to insecticides were observed in *Ae. aegypti formosus* and *Ae. aegypti aegypti* (Moore et al. 2013). This highlights the epidemiological significance of the various strains of this species, underscoring the need for thorough characterization to enhance vector control strategies.

In our recent study, we identified a morphological variant, known as *Ae. aegypti var. luciensis* form various locations within India (Kumar et al. 2022). This variant has previously been observed in India, Anguilla and Lagos, where it was initially found in tree holes (Connal 1927; Parker et al. 1983; Kumar et al. 2022). However, in our recent surveys, we identified both the morphotypes coexisting in the breeding places, indicating recent expansion and adaptation of this mosquito variant. The genetic differences between the two morphotypes are evident from our previous crossing experiments (Kumar et al. 2022). These experiments demonstrated that crosses between the two morphotypes produced mixed offspring, but each cross developed into the same type as the parents. This finding reinforces the notion that the observed morphological differences are underpinned by genetic factors rather than being solely ecologically controlled. Although genetic analysis of the COI and ITS2 regions did not reveal clear genetic differences, likely due to a lack of reproductive isolation and the potential for genome mixing from inbreeding. Genome-level comparisons could provide detailed insights on the closely related organisms.

In this study, we conducted genome-level comparison of these two morphotypes. Our analysis indicated significant genetic differences between the two subtypes; however, specific regions or genes could not be identified. Nonetheless, these differences suggest that there are genetic variations that may be influencing their demographic histories, current population dynamics, and other biological aspects.

## Methods and materials

### Mosquito collection and genomic DNA extraction

*Ae. aegypti* morphotypes were collected from Madurai district of Tamil Nadu, India. The sample collection methodology is described in our previous studies (Kumar et al. 2021, 2022; Sumitha et al. 2023) *Ae. aegypti* mosquitoes were identified following the taxonomic keys (Theobald 1901; Christophers 1933, 1960). Nine samples included four type forms (hereafter called ‘normal’) and its morphological variants (hereafter called ‘variant’) were used for sequencing. Genomic DNA extraction was carried out using commercially available DNeasy® Blood & Tissue Kit (Qiagen, USA). DNA quantity and quality was checked using the Nanodrop (Thermo Scientific), and Qubit fluorometer.

### Illumina library preparation and genome sequencing

Approximately 10 ng of DNA from each sample was fragmented, end-repaired and A-tailed using QIASeq FX DNA Library Preparation protocol (Cat#180475). The library was generated by ligating the adenylated fragments with index-incorporated Illumina adapters. The library underwent 10 cycles of Indexing-PCR and the amplified fragments were purified with Sera-MagTM Select beads (Cytiva,#29343057) and quantification was done using Qubit. The fragment size distribution was analyzed using Agilent 2200 TapeStation (Salowsky et al. 2013). The sequencing was done on Illumina NovaSeq 6000 (Illumina, San Diego, USA) using 150 bp paired-end chemistry. The reads obtained after sequencing run were de-multiplexed using Bcl2fastq2 v2.20 and FastQ files were generated. The library construction and sequencing were done at Genotypic technologies, Bengaluru.

### Data Analysis

The raw Illumina pair-end reads were subjected to quality check and the adapter sequences were trimmed-off using Trimgalore-v0.4.04 tool (Krueger 2015). Low quality reads with Phred value (Q) less than 30 were removed from downstream analysis. The high-quality reads were mapped against reference *Ae. aegypti* genome (*Aedes aegypti AaegL5*.*0*) using BWA-v0.7.55 aligner (Li and Durbin 2009). Alignment data was further analysed using Samtools-v1.96 (Li et al. 2009). The PCR duplicate reads were removed from aligned data using PICARD-v1.1027 tool (https://broadinstitute.github.io/picard/). The variants in each sample were identified using GATK-v4.2.6.18 HaplotypeCaller (McKenna et al. 2010). GATK is capable of calling SNPs and indels simultaneously via local assembly of haplotypes in the genomic region. Finally, the annotation was carried out for all the identified variants using SnpEff-v3.3h9 (Cingolani et al. 2012). The identified variants were further used for phylogenetic analysis using vcf2pop software (Subramanian et al. 2019).

## Results and discussion

### Read mapping and alignment to the reference genome

Around 27 to 46 million reads were obtained from nine mosquitoes with sequencing coverage in the range of 7 to 11X. One of the variant samples, M7, had comparatively low read counts and the lowest coverage (Table 1); thus, it was excluded from further analysis. Consequently, the dataset included eight samples: four variants (M1, M6, M8, M9) and four normals (M2 - M5). The reads were subjected to quality control and the reads with Q>30 and length >20 bases were retained from further analysis. This stringent filtering ensured that more than 99% of the reads in all samples met these criteria. These high-quality reads were then mapped to the reference mosquito genome. The percentage of mapped reads varied from 97.55% to 98.92% across the eight samples, indicating a high level of alignment accuracy and consistency with the reference genome.

**Table 1.**
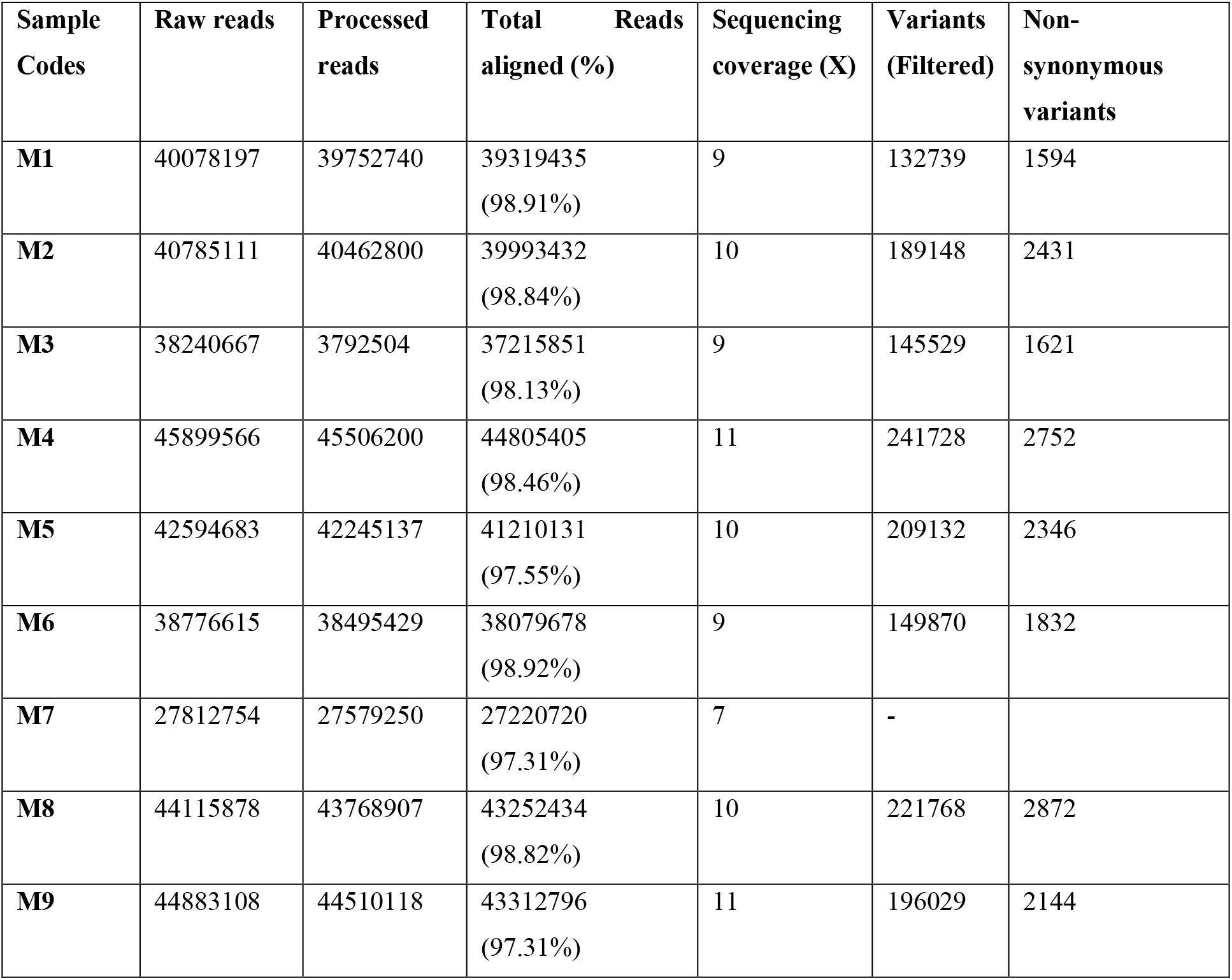
Details on the number of raw reads, processed reads, total variants, and non-synonymous variants obtained from each sample.

| **Sample Codes** | **Raw reads** | **Processed reads** | **Total Reads aligned (%)** | **Sequencing coverage (X)** | **Variants (Filtered)** | **Non-synonymous variants** |
| --- | --- | --- | --- | --- | --- | --- |
| **M1** | 40078197 | 39752740 | 39319435<br>(98.91%) | 9 | 132739 | 1594 |
| **M2** | 40785111 | 40462800 | 39993432<br>(98.84%) | 10 | 189148 | 2431 |
| **M3** | 38240667 | 3792504 | 37215851<br>(98.13%) | 9 | 145529 | 1621 |
| **M4** | 45899566 | 45506200 | 44805405<br>(98.46%) | 11 | 241728 | 2752 |
| **M5** | 42594683 | 42245137 | 41210131<br>(97.55%) | 10 | 209132 | 2346 |
| **M6** | 38776615 | 38495429 | 38079678<br>(98.92%) | 9 | 149870 | 1832 |
| **M7** | 27812754 | 27579250 | 27220720<br>(97.31%) | 7 | - |  |
| **M8** | 44115878 | 43768907 | 43252434<br>(98.82%) | 10 | 221768 | 2872 |
| **M9** | 44883108 | 44510118 | 43312796<br>(97.31%) | 11 | 196029 | 2144 |

Although depth of sequencing data is relatively low (9-11X), the high quality of the reads and the high percentage of mapped reads provide confidence in our findings. Low coverage data has been effectively used in various comparative genomic studies to uncover significant genetic differences and evolutionary insights. For instance, similar approaches have been utilized in studies on *Anopheles gambiae* for malaria vector genomics (Dennis et al. 2024), *Ae. aegypti* for population structure analysis (Lee et al. 2019; Gómez-Palacio et al. 2024), green lacewings (Chrysopidae) to study their phylogeny (Wang et al. 2021).

### Identification of the variants and comparison of the morphotypes

The aligned data from all the samples were processed to remove PCR duplicate reads before proceeding for calling the variants. The variants were filtered using standard parameters such as minimum read depth of 20 and base quality of 30. A total 132,739 variants in M1 (variant type) to 241,728 in M4 (normal type) were identified on comparing with the reference genome (Table 1). However, the number of non-synonymous variants ranged from 1,594 to 2,872.

To ensure robustness, we combined all four samples from each type and retained the variants present in all samples, resulting in 7,181 variants in the normal type and 4,513 variants in the variant type. The overall high level of genetic diversity in the normal samples compared to the variant samples is possible due to a larger population size and greater adaptability. This species is highly invasive and exhibits significant genetic diversity that may enhance its ability to adapt to varying environmental conditions (Birader 2023; Zhao et al. 2023). In contrast, the low level of diversity observed in variant samples suggests that this variant may have a more restricted genetic pool due to recent divergence. However, this cannot be conclusively stated due to a lack of data on morphological variants. Another possible reason for the lower genetic diversity is that since the two types coexist, they may face survival competition in the field, with the one exhibiting greater adaptability potentially gaining an advantage. This situation warrants further investigation through field entomological studies and assessments of the life history parameters of both morphotypes.

For further comparison of the morphotypes, we combined variants form all the eight samples that resulted in 10,514, including; 9840 SNPs and 260 insertions and 414 deletions. Of the total, 10031 variants were chromosomal, while 483 were found in scaffolds. The frequency of variants identified across the three chromosomes is shown in Table 2. The number of variants were in similar range across the chromosomes in both morphotypes. The transition/transversion (Ts/Tv) ratios were also comparable between the two types with Ts/Tv ratios of 1.46 and 1.45, respectively indicating similar mutation processes are occurring in both morphotypes.

**Table 2.** Chromosome-wise distribution of variants for both the normal type and the variant type of *Aedes aegypti*.

|  | **Variants (SNP + InDels)** |  |
| --- | --- | --- |
|  | **Normal** | **Variant** |
| **Chromosome 1** | 2268 | 1298 |
| **Chromosome 2** | 2291 | 1456 |
| **Chromosome 3** | 2343 | 1453 |

On comparing the two types, we found that only 1,180 variants were common between them. The normal samples exhibited 5,995 unique variants, while the variant type had 3,327 unique variants. Clustering analysis revealed two distinct clusters, indicating clear genetic distinctions between the two types (Figure 1). This divergence suggests that each morphotype has developed a unique genetic signature, despite some overlap. However, these differences highlight the need for recognizing these morphotypes for epidemiological understandings as well as for developing effective vector control measures.

**Figure 1:**
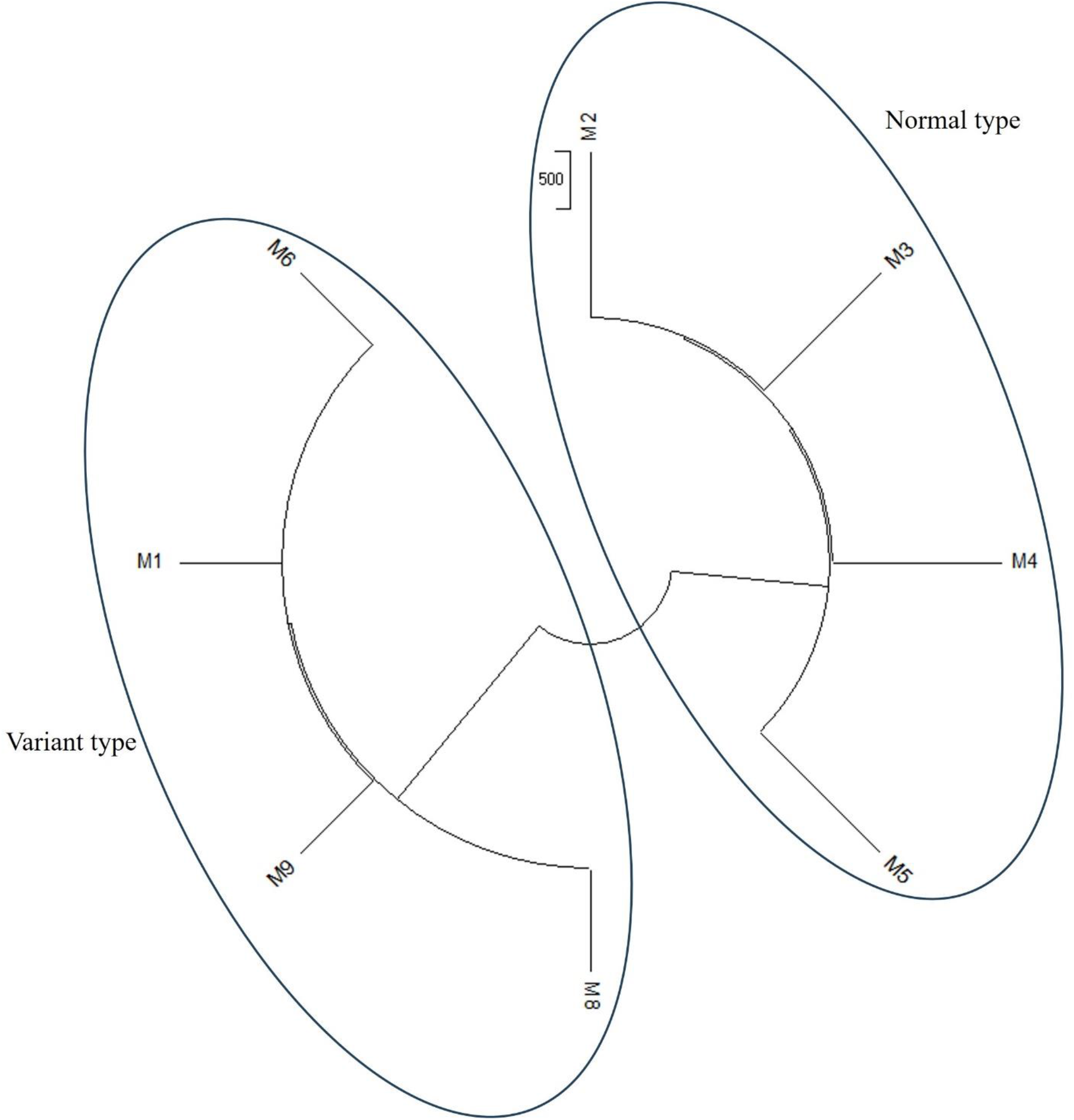
Neighbor-joining tree depicting genomic differences between two morphotypes of *Aedes aegypti*.

### Characterization of the unique and common variants among morphotypes

The annotation of 10,514 variants showed 30,100 effects. Each variant can have multiple effects depending on its genomic context, such as affecting several transcripts of a gene or impacting overlapping or closely situated genes (Shen et al. 1999; Lutz et al. 2019; Bohry et al. 2021). Variants in regions with alternative splicing can influence multiple transcript variants, increasing the number of effects (Pagani and Baralle 2004; Palaniswamy et al. 2010; Monlong et al. 2014; Hsiao et al. 2016; Smith et al. 2018; Joshi et al. 2021). However, the majority of the effects were in non-coding regions, with only 139 found in exons, including 82 missense mutations and four high-impact mutations. Of the high-impact variants, three resulted in frameshift mutations and one led to a splice acceptor mutation. Two of these high-impact mutations were found in the AAEL024218 gene (uncharacterized), which also showed multiple missense mutations. The other two high-impact mutations were observed in AAEL021592 and AAEL020899. Missense mutations were identified in 18 genes, as shown in Table 3.

**Table 3.** List of genes in which missense mutations have been identified.

| **Sl. No.** | **Gene ID** |
| --- | --- |
| 1 | AAEL004426 |
| 2 | AAEL004546 |
| 3 | AAEL019423 |
| 4 | AAEL019736 |
| 5 | AAEL020899 |
| 6 | AAEL021592 |
| 7 | AAEL022027 |
| 8 | AAEL023871 |
| 9 | AAEL024218 |
| 10 | AAEL024457 |
| 11 | AAEL024777 |
| 12 | AAEL025031 |
| 13 | AAEL025942 |
| 14 | AAEL026638 |
| 15 | AAEL027214 |
| 16 | AAEL027513 |
| 17 | AAEL027850 |
| 18 | AAEL021293 |

Among the unique variants in the normal samples, 186 were homozygous for alternate alleles, indicating that those positions might be completely different from the normal samples. Similarly, 318 variants in the normal samples were monomorphic but differed from those in the variant samples. Analysis of the genes closer to these unique non-coding variants using the DAVID system (Sherman et al. 2022) indicated that the most enriched terms included synapse organization, structure morphogenesis, cell-cell adhesion, transcription regulation G-protein coupled receptor (GPCR) pathway, voltage gated channels. Although the majority of these variants were located in non-coding regions. However, the existence of the unique variants in each types indicates the different demographic histories or the competitive dynamics between the two morphotypes. Future studies should focus on the ecological interactions and competitive relationships between these morphotypes to better understand the factors driving their genetic divergence. The differences between the two types might also be associated with variations in vector competence, or other biological aspects which could have significant implications for their survival and disease transmission.

### Conclusion

In conclusion, our findings highlight the significant genetic differentiation between the normal and variant morphotypes, despite their lack of reproductive isolation. The higher genetic diversity in the normal type suggests a broader adaptive potential, while the variant type may be constrained by competitive pressures. The observed low diversity in the coding regions might be attributed to the low coverage of the data. It is also possible that the genetic closeness of the two morphotypes has resulted in limited differentiation. Both morphotypes coexist in the same breeding containers; however, historically, the variant was identified in rock-holes (Parker et al. 1983). These differences in demographic history and population dynamics may have impacted the level of coding and non-coding differentiation between the two morphotypes. Future research should aim to unravel the ecological interactions, a comprehensive understanding of their evolutionary trajectories and the susceptibility to different arboviruses. This will help to clarify the factors driving these differences.

## Acknowledgements

We would like to thank ICMR for intramural support. We would like to express our sincere gratitude to Dr. Manju Rahi, Director, ICMR-VCRC, Puducherry for her invaluable support. G Navaneetha Pandiyan and Melveettil Kishor Sumitha would like to thank Madurai Kamaraj University for supporting their research.

## Statements and declarations

The authors state that none of the work described in this publication appears to have been influenced by any known competing financial interests or personal ties.

## CRediT authorship contribution statement

**Bhavna Gupta:** Conceptualization, Formal analysis, Funding acquisition, Methodology, Project administration, Supervision, Visualization, Writing – original draft. **G Navaneetha Pandiyan:** Data curation, Writing – review and editing. **Melveettil Kishor Sumitha:** Data curation, Writing – review and editing. **Rajaiah Paramasivan:** Resources. **Mariapillai Kalimuthu:** Data curation, Writing – review and editing.

